# D_1_, not D_2_, dopamine receptor activation dramatically improves MPTP-induced parkinsonism unresponsive to levodopa

**DOI:** 10.1101/2020.09.20.305375

**Authors:** Richard B. Mailman, Yang Yang, Xuemei Huang

**Affiliations:** Departments of Pharmacology and Neurology, Penn State University College of Medicine, Hershey PA 17033

**Keywords:** MPTP, parkinsonism, Parkinson’s disease, dopamine, dopamine D_1_ receptors, dopamine D_1_ agonists, dihydrexidine

## Abstract

Levodopa is the Parkinson’s disease standard-of-care, but continued loss of dopamine neurons with disease progression decreases its bioconversion to dopamine, leading to increased side effects and decreased efficacy. In theory, dopamine agonists could equal levodopa, but no approved oral “dopamine agonist” matches the efficacy of levodopa. Although there are consistent data in both primate models and in Parkinson’s disease showing that selective high intrinsic activity D_1_ agonists can equal levodopa, there are no data on whether such compounds would be effective in severe disease when levodopa efficacy is lower or even absent. We compared two approved antiparkinson drugs (levodopa and the D_2/3_ agonist bromocriptine) with the experimental selective D_1_ full agonist dihydrexidine in two severely parkinsonian MPTP-treated non-human primates. Bromocriptine caused no discernable improvement in parkinsonian signs, whereas levodopa caused a small transient improvement in one of the two subjects. Conversely, the full D_1_ agonist dihydrexidine caused a dramatic improvement in both subjects, decreasing parkinsonian signs by ca. 75%. No attenuation of dihydrexidine effects was observed when the two subjects were pretreated with the D_2_ antagonist remoxipride. These data provide evidence that selective D_1_ agonists may provide profound antiparkinson symptomatic relief even when the degree of nigrostriatal degeneration is so severe that current drugs are ineffective. Until effective disease-modifying therapies are discovered, high intrinsic activity D_1_ agonists may offer a major therapeutic advance in improving the quality of life, and potentially the longevity, of late stage Parkinson’s patients.

## 1. Introduction

The motor signs of Parkinson’s disease (PD) (Parkinson, 1817) are a result of dopamine deficiency due to progressive degeneration of dopamine neurons cells in the substantia nigra pars compacta (Ehringer and Hornykiewicz, 1960; Hornykiewicz, 1963). Despite recent insights into the biology of PD, no preventative or restorative therapy is yet available, thus the gold-standard therapy remains levodopa (Cotzias et al., 1969), a dopamine precursor that dramatically increases the dopamine levels in the brain (Davidson et al., 1971). Bioconversion of levodopa *in situ* depends on the residual dopamine terminals that are lost with disease progression, causing a decrease in efficacy and an increase in side effects such as wearing off and freezing (Connolly and Lang, 2014). Ultimately, a patient will often be wheelchair- or bed-bound if they do not succumb to injury, infection, or another disorder.

Levodopa indirectly activates dopamine receptors comprising six G protein-coupled receptors in two pharmacological groups: D_1_–like (D_1_ and D_5_); and D_2_-like (D_2L,_ D_2S_, D_3_, D_4_) (Neve and Neve, 1997). In theory, a dopamine agonist should be able to equal or exceed the efficacy of levodopa by targeting one or more of the essential dopamine receptors, yet all approved “dopamine agonists” (selective for D_2_-like receptors) are less efficacious than levodopa, especially in late stages of PD (Mailman and Huang, 2007; Rothman et al., 2000). There are data, however, suggesting a critical role of D_1_-like receptors, specifically the D_1_. Although data with the partial agonist SKF38393 (Setler et al., 1978) showed little efficacy (Boyce et al., 1990; Close et al., 1985), the first full D_1_ agonist dihydrexidine (Brewster et al., 1990; Lovenberg et al., 1989; Mottola et al., 1992) caused profound antiparkinson effects in MPTP-treated African green monkeys, essentially eliminating all parkinsonian signs acutely (Taylor et al., 1991). Later full D_1_ agonists, such as A-77636 (Kebabian et al., 1992) and A-86929, a structural and pharmacological cousin of dihydrexidine (Michaelides et al., 1995)], also produced profound antiparkinson effects (Shiosaki et al., 1996). Importantly, ABT-431 (an A-86929 prodrug), was equi-efficacious to levodopa in two clinical trials (Rascol et al., 1999; Rascol et al., 2001).

Levodopa efficacy decreases as dopamine terminals are lost with PD progression (Lewis et al., 2011; Lewis et al., 2007), thus selective activation of D_1_ receptors might provide profound symptomatic relief even in late-stage PD patients who are unresponsive to levodopa. Goulet and Madras (2000) reported that D_1_ agonists (including dihydrexidine) were more effective in severe than in moderate parkinsonian MPTP-treated cynomologous monkeys. Consistent with Goulet and Madras (2000), we now report data from two MPTP-treated African green monkeys with severe MPTP-induced parkinsonism in which dihydrexidine caused profound antiparkinson effects that were greater than seen with either levodopa or the D_2_ agonist bromocriptine. Moreover, these effects were not attenuated by prior treatment with a D_2_ antagonist. These data suggest that pharmacological intervention with D_1_ agonists may have therapeutic value in advanced PD where no other therapy is of marked utility.

## 2. Materials & Methods

### 2.1 Materials

Dihydrexidine was synthesized as described previously (Brewster et al., 1990; Ghosh et al., 1996). Bromocriptine was a gift from Novartis (formerly Sandoz Pharmaceuticals, East Hanover, NJ), and remoxipride was a gift from AstraZeneca (formerly Astra AB, Södertälje Sweden). Levodopa and benserazide were purchased commercially.

### 2.2 MPTP treatment and animal care

A group of adult male monkeys (*Cercopithecus aethiops sabaeus*) from St. Kitts, West Indies were injected intramuscularly with 0.3 to 0.4 mg/kg MPTP given four or five times over a five-day period. They were housed individually in standard primate cages in natural daylight. Access to food and water was unlimited. Care and treatment of these monkeys were in compliance with the US Public Health Service Guide for the Care and Use of Animals (1985), and the protocols were approved by the by the Axion/St. Kitts Biomedical Research Foundation committee. All experimental monkeys were carefully monitored for their general level of function, motor coordination, and oral intake before and after MPTP treatment, and any monkey failing to take established minimums of fluid or food was assisted as necessary to ensure adequate nutrition and hydration. Daily physiotherapy consisting of regular passive range of motion exercises and monitoring of temperature and respiration rates was done based on severity of impairment. Medical complications of severe parkinsonism, when they occur, were evaluated and treated as necessary to minimize the impact of non-parkinsonian disease or disability on the function of the animal. Details of the handling of these animals have been published previously (Taylor et al., 1994; Taylor et al., 1997).

Of this group, several animals became unusually parkinsonian, and were no longer able to sit upright or to feed themselves. When the excess parkinsonism was noted, these subjects received special care that included special padded bedding to minimize contact with the hard cage surfaces, and feeding assistance and special nursing care that included may feeding by humans of a softened commercial monkey chow via a large syringe. Of these animals, some were used for assessment with nigral fetal tissue grafts into the striatum and two monkeys were selected for this pharmacological rescue study reported here. After these animals failed to improve spontaneously over several weeks, levodopa was tried as a rescue medication, but there was no adequate response in either subject. The recommendation from the attending veterinarian was to euthanize the subjects. It was recognized, however, that these two subjects represented a model of end-stages of Parkinson’s disease in which patients are bed-ridden, can no longer self-feed, and for which there is no currently available effective therapy. Based on data showing the large effect size of dopamine D_1_ full agonists in the MPTP model of moderate PD, it was felt that these subjects provided the unusual opportunity to determine if a D_1_ agonist might offer hope in advanced PD. Thus, a controlled cross-over acute comparison of current standards of care (levodopa and D_2_ agonists) with dihydrexidine, a full D_1_ agonist, was felt to be warranted. Because only two subjects were available, no statistical analysis was possible, but each subject received each of the four treatments, and the videotapes were rated by two trained observers who were blinded to treatments. The ratings of the videotapes in Figures 2 and 3 are complete and unedited.

Prior studies in this model have shown that severe animals of the type used in this study have Parkinson scores of ca. 60-70, and unlike moderate or mildly parkinsonian animals (scores of 8-17) do not recover with time (Elsworth et al., 2000). Such severe animals have an overall dopamine depletion in the striatum of >99%, whereas moderate animals, such as used in our earlier study of dihydrexidine (Taylor et al., 1991) have depletions of 90-95%.

### 2.3 Drug administration

There was a one-day interval between each drug trial. The two subjects were tested on the same day. The order of treatment was levodopa, bromocriptine, dihydrexidine, and dihydrexidine plus remoxipride.

### 2.4 Videotaping and blind ratings

The subjects were videotaped, and the effects rated by two trained observers who were blinded to the order of interventions. These observers had achieved a coefficient of concordance (Kendall’s) greater than 0.95 on all behaviors. The ratings used an observation scale that has been reported in detail (Taylor et al., 1994; Taylor et al., 1997), the components of which are summarized in Figure 1.

**Figure 1.**
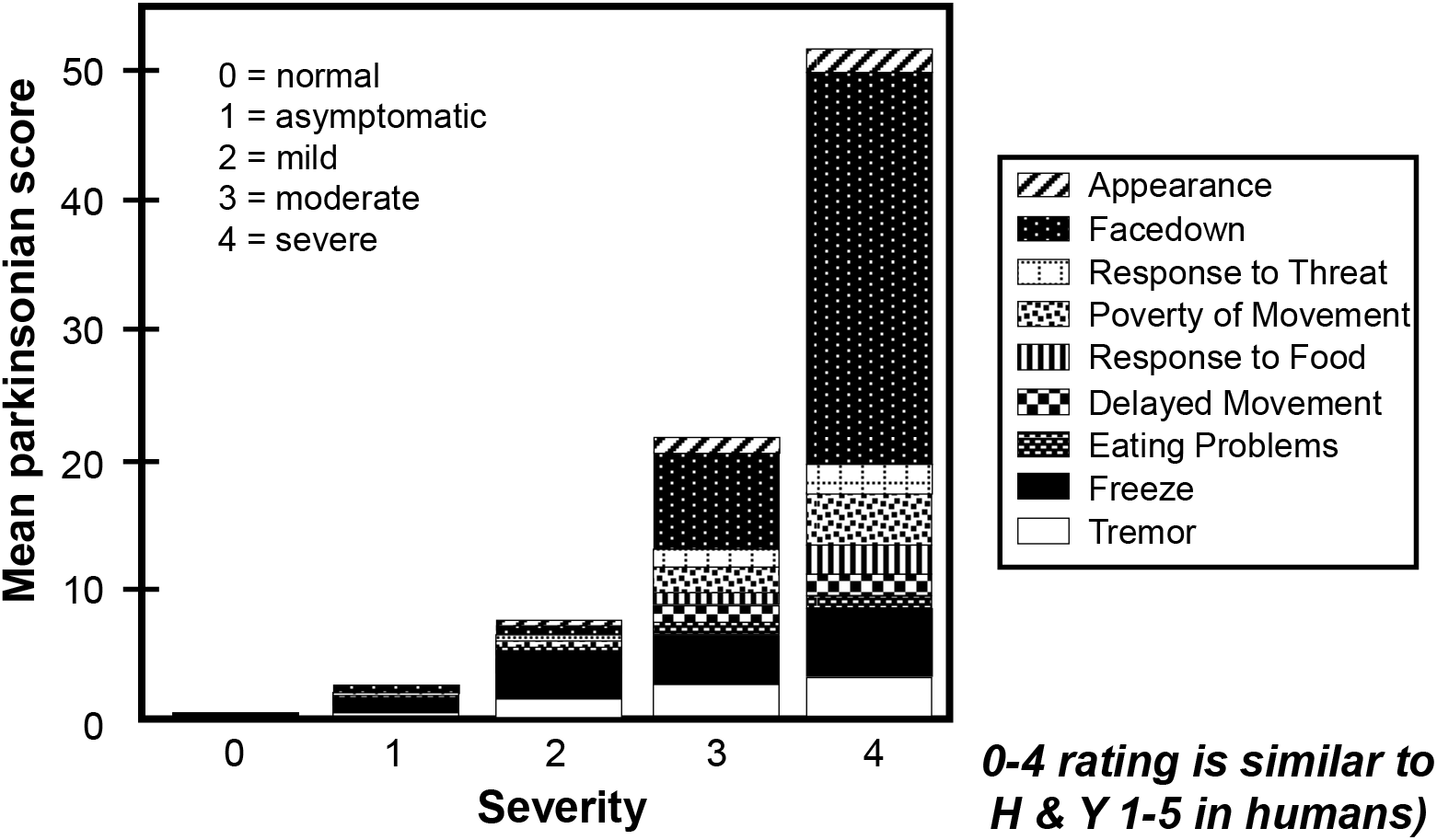
Primate rating scale. Scale for rating of parkinsonian signs in these subjects [from(Taylor et al., 1994; Taylor et al., 1997)]

## 3. Results

Although only two severely parkinsonian subjects were available, the study was felt to be feasible because prior research in moderately parkinsonian MPTP-treated non-human primates (NHPs) has shown a very large effect size of the full D_1_ agonist dihydrexidine (Taylor et al., 1991). Moreover, because dihydrexidine and levodopa have relatively rapid metabolism, carry-over effects are minimal, and an inter-subject design could be used with each subject serving as its own control. We thus used a blinded cross-over design in which each subject was treated in separate experiments with: 1) the D_1_ agonist dihydrexidine; 2) the D_2_ agonist bromocriptine; 3) levodopa; and 4) the D_2_ antagonist remoxipride plus dihydrexidine. All of the trials were videotaped, and scored blindly by two experienced and inter-reliable raters (see Supplementary Material).

Although levodopa is known to cause a robust response in moderately parkinsonian MPTP-NHPs, prior uncontrolled rescue studies in these two subjects showed minimal response. This was confirmed by the relative lack of response to levodopa plus benserazide in subject 236, and only a modest response in subject 201 (Figure 2-left panel). When challenged with the D_2_ agonist bromocriptine, neither subject showed any noticeable improvement (Figure 2, right panel).

**Figure 2.**
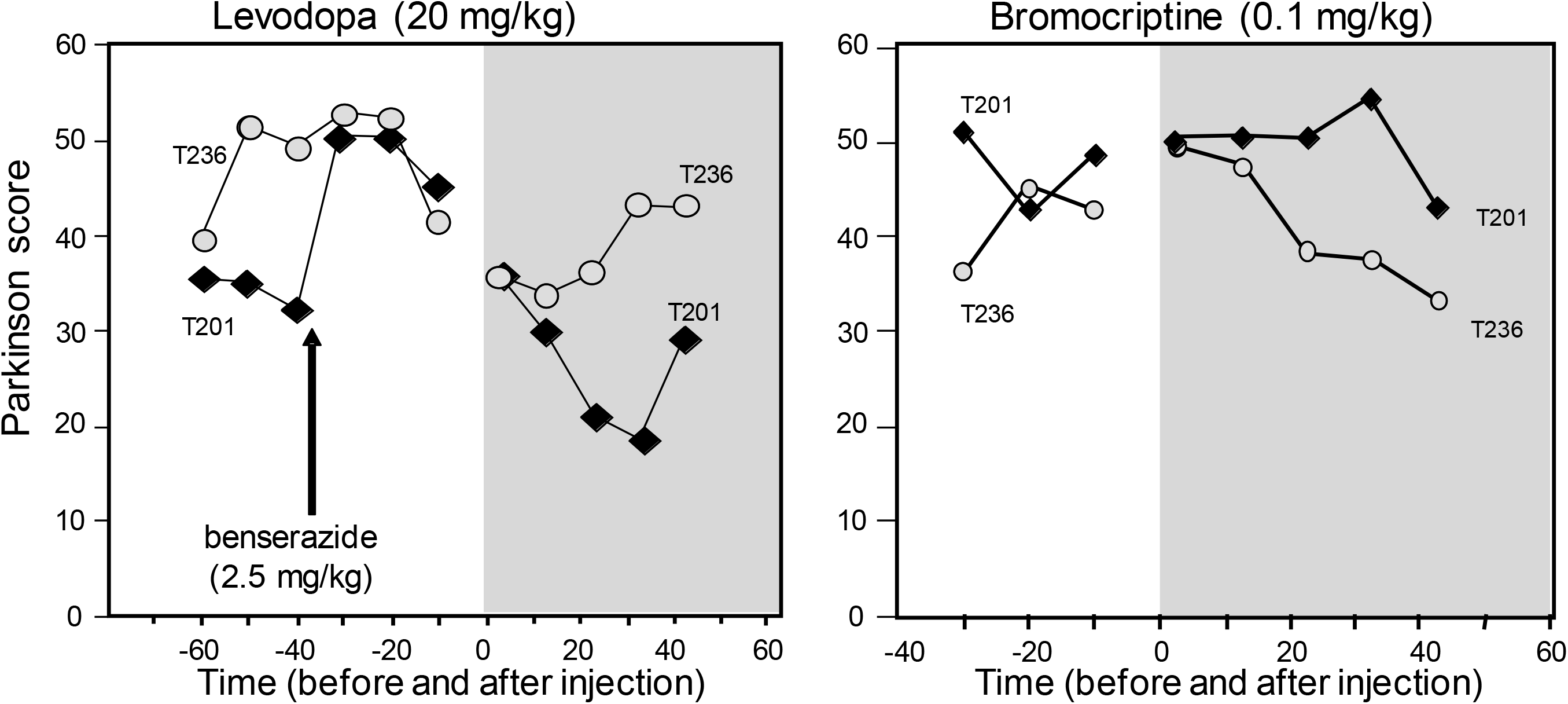
(Left) Time course of effects of levodopa in two MPTP-treated NHPs (T236 and T201) after injection of benserazide, a decarboxylase inhibitor (arrow indicates injection of benserazide). (Right). Time course of effects of the D_2_ agonist, bromocriptine. Shaded area denotes time after injection of active drug (i.e., levodopa in left panel; bromocriptine-right panel). Results are the average of the scores of two raters.

Conversely, when treated with 0.6 mg/kg of dihydrexidine, both subjects showed a remarkable response. Prior to treatment, they could neither sit, stand, or feed themselves, they were moving significantly and taking food (Figure 3, left panel). Because dihydrexidine is only ten-fold D_1_:D_2_ selective, there was the possibility that the D_1_ effects might require concomitant D_2_ activation, or indeed, the D_2_ activation was solely responsible for the observed effects. Thus, the animals were pretreated with the D_2_ antagonist remoxipride, and then challenged again with dihydrexidine. Even with the pretreatment with remoxipride, dihydrexidine again caused a marked decrease in parkinsonian signs in both subjects (Figure 3, right panel).

**Figure 3 (Left).**
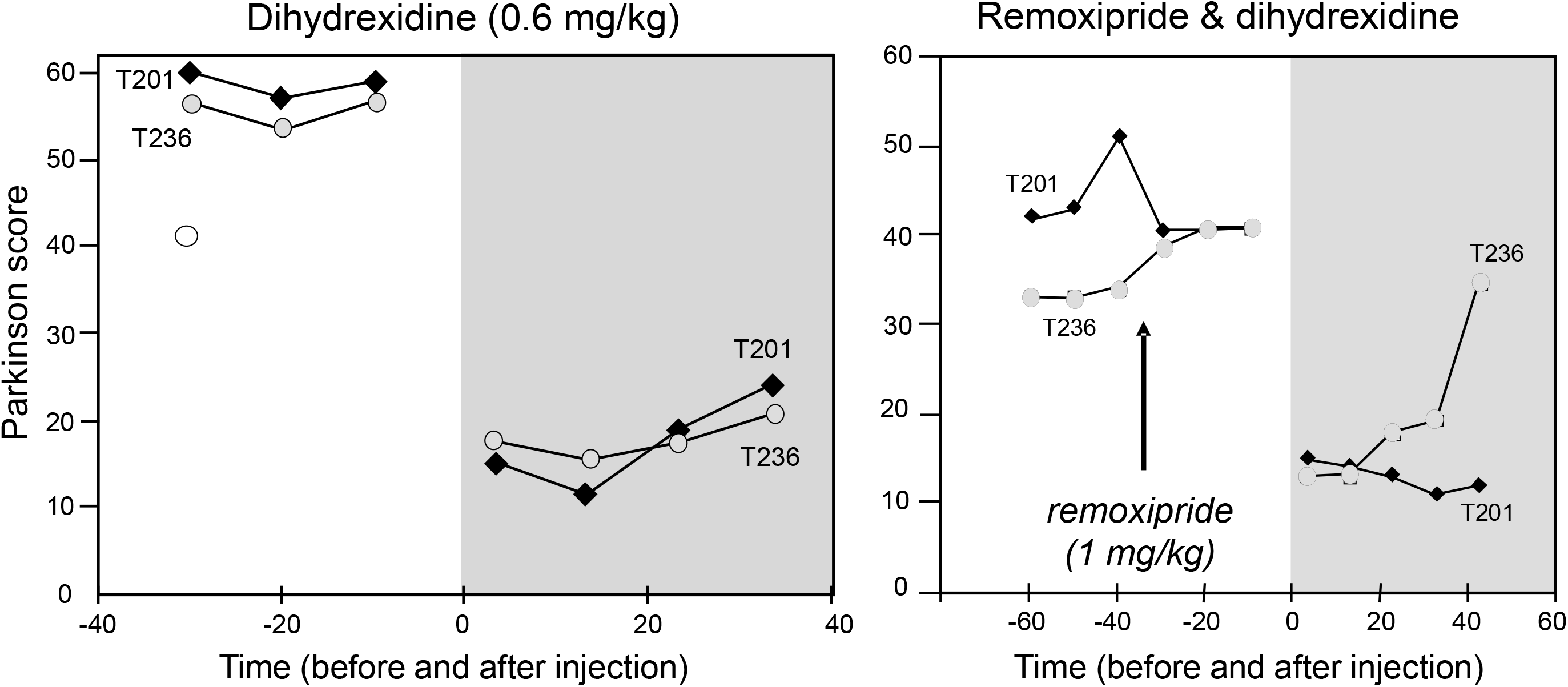
Effects of the full D_1_ agonist dihydrexidine on parkinsonian signs in two MPTP-treated NHPs (subjects T236 and T201). (Right) Effects of pretreatment with D_2_ antagonist remoxipride (arrow indicates the time of remoxipride injection) on subsequent treatment with dihydrexidine. Shaded area denotes time after injection of dihydrexidine.

## 4. Discussion

### 4.1 Profound antiparkinson effects of a D_1_ full agonist

We had previously reported in moderately parkinsonian MPTP-treated African green monkeys that dihydrexidine essentially eliminated all parkinsonian signs (Taylor et al., 1991). The current data show that there was also a very large effect size in severely parkinsonian subjects of the same species. These data are in partial agreement with Goulet and Madras (2000) who reported that two D_1_ agonists (dihydrexidine and SKF81397) both had significant, but not dramatic, effectiveness in four severely affected MPTP-treated cynomologous monkeys. Conversely, they saw no major effects in “mild” subjects, contrary to our earlier report showing consistent dramatic effects in nine subjects (Taylor et al., 1991). Goulet and Madras (2000) also reported a large effect size of two D_2_-like agonists (PHNO and quinelorane), albeit with side effects, whereas we saw no effect with the D_2_-like agonist bromocriptine. Their study did not compare these dopamine agonists with levodopa, whereas ours showed that there was no effect of a large dose of levodopa. Another difference was that we saw very large effects at 0.6 mg/kg of dihydrexidine (consistent with doses to get maximal effects in rat models), whereas they required doses five times higher. Because of the practical and ethical difficulties in working with severely parkinsonian non-human primates, both studies had small numbers of subjects, and used animals of different species and with different scoring systems. Nonetheless, despite these limitations, our results are consistent with what is seen in very advanced PD patients where levodopa effects are lost and D_2_ agonists are not effective.

### 4.2 Why should D_1_ agonists be effective?

There is a sound neurobiological, as well as pharmacological, underpinning for these findings. Models of basal ganglia neurocircuitry in both the normal and pathophysiologic state in PD (Albin et al., 1989; DeLong, 1990) posit that coordinated movement is regulated by two parallel and segregated pathways (direct and indirect) in the basal ganglia that are formed by the GABAergic medium spiny projection neurons of the striatum that enables movement via disinhibition of thalamo-cortico circuits. Early studies showed that the striatum had the highest expression of D_1_ receptors (Schulz et al., 1985), and D_1_ antagonism could block dopaminergic motor stimulation (Mailman et al., 1984), suggesting that the D_1_ receptor-mediated direct pathway might facilitate movement stimulatory actions, whereas the D_2_ receptor-mediated indirect pathway is inhibitory (Gerfen et al., 1990). The dopamine deprivation in PD impairs the disinhibition of thalamo-cortico circuits through these two distinct pathways, causing a general hypokinetic state. It has, however, been argued that colocalization of D_1_/D_2_ receptors in a minor proportion of striatal neurons, and even D_3_ and D_5_ receptors, also may be important (Surmeier et al., 1992). The current data show that, at least in severe parkinsonism, activation of only D_1_ is entirely adequate for marked antiparkinson effects, contrary to a long-held dogma (Cedarbaum and Schleifer, 1990).

### 4.3 Strengths and limitations of the MPTP-model

The impact of our findings on future clinical results depends on the ability of our findings to translate to clinical PD. Although the MPTP-model has not led to translatable breakthroughs in understanding PD etiology, symptomatically there has been excellent correlation of the motor efficacy of drugs used in PD with its motor improvement in MPTP-treated NHPs. This is not surprising because there is major loss of striatal dopamine neurotransmission in both PD and in MPTP-NHP models. This suggests a higher likelihood of our findings translating to humans then when the MPTP-model is used for disease-etiology purposes. Indeed, our report showing profound efficacy of dihydrexidine in moderately parkinsonian MPTP-NHPs (Taylor et al., 1991) predicted that ABT-431 would be clinically effective because its active species A68929 is structurally and pharmacologically similar dihydrexidine (Michaelides et al., 1995). This was confirmed in studies in moderate PD that showed clinical efficacy essentially equal to levodopa (Rascol et al., 1999; Rascol et al., 2001) that cannot be achieved with currently approved oral dopamine agonists. One important potential limitation relates to disease progression. During the course of PD, there are complex pathological changes that unfold, and these differ within subjects as well as within similar parkinsonian disorders (Du et al., 2018; Lewis et al., 2018). Even though striatal D_1_ receptors are essentially all post-synaptic to degenerating dopamine terminals, and even though they are present post-mortem, neither their functional state, nor that of the circuitry mediated by the direct pathway, have been shown to retain near-normal status. Whether this is a concern awaits both experimental and clinical studies.

### 4.4 Intrinsic pharmacological issues

The original selective full D_1_ agonists had notable pharmaceutical limitations (Rascol et al., 1999; Rascol et al., 2001; Taylor et al., 1991), and also potential dose-limiting side effects (Blanchet et al., 1998). This has led to the widespread feeling that the D_1_ receptor was not a druggable target, although others felt this was simply because adequate efforts had not been made (Mailman et al., 2001; Mailman and Huang, 2007). Recently, a new series of D_1_ agonists was discovered that overcome the pharmaceutical limitations of earlier drugs (Brodney et al., 2014; Davoren et al., 2014), and which have been shown to be safe and have antiparkinson effects in mid-stage disease (Papapetropoulos et al., 2018; Sohur et al., 2018). Many of the earlier D_1_ selective agonists like SKF38393, but very low intrinsic activity (Setler et al., 1978), possibly explaining why it failed in both MPTP-NHP and human trials (Boyce et al., 1990; Close et al., 1985) unlike later full agonists (Kebabian et al., 1992; Lovenberg et al., 1989; Michaelides et al., 1995; Montastruc et al., 1999; Rascol et al., 2001; Shiosaki et al., 1996; Taylor et al., 1991).

More recently, it has been recognized that drugs may cause differential activation of signaling pathways modulated by a single receptor, a mechanism termed functional selectivity or biased signaling (Urban et al., 2007). There are numerous studies showing that D_1_ selective ligands can evoke this mechanism, although there is some controversy about its nature (Lee et al., 2014). The specific signaling properties may, therefore, influence the clinical effects. Although we believe dihydrexidine is an unbiased full D_1_ agonist, *in vitro* studies of Yano et al. (2018) suggested it was a full agonist when binding to D_1_ Gα_S_, but only a partial agonist for D_1_ Gα_OLF_ (the latter G protein critical for D_1_ actions in the striatum). On the other hand, we have reported that dihydrexidine is a full agonist for D_1_-adenylate cyclase activation in monkey, human, rat, and mouse striatum (Gilmore et al., 1995; Lovenberg et al., 1989; Mottola et al., 1992; Watts et al., 1993; Watts et al., 1995), whereas the results of Yano et al. (2018) would require it to be a partial agonist. Moreover, in Gα_OLF_^+/−^ mice, the efficacy of both dopamine and dihydrexidine was identical, half that of wild-type mice (unpublished data). Although the *in vivo* data strongly support the hypothesis that dihydrexidine is a full agonist at Gα_OLF_ mediated signaling, the question may be worthy of additional study, especially as a highly-biased D_1_ partial agonist recently was reported to be effective in PD (Papapetropoulos et al., 2018). Interestingly, Interestingly, thee new clinical D_1_ agonist candidates are not all full agonists, but are functionally selective with no D_1_ mediated β-arrestin2 recruiting activity (Gray et al., 2018).

### 4.5 Conclusions

Despite its limitations, our data suggest that D_1_ agonists may be very useful at stages of Parkinson’s disease that are now essentially untreatable. Because very advanced Parkinson’s patients have no available treatment, they suffer from many secondary effects of the disease resulting from inactivity, falls, swallowing difficulty, and many non-motor symptoms. For these reasons, to our knowledge there has never been an interventional study in patients in this very advanced state, in part because there was no potential therapy that justified such clinical trials. Based on the results of this study and influenced by that of Goulet and Madras (2000), using the oral partial D_1_ agonist PF-06412562 (Papapetropoulos et al., 2018), we have recently concluded what we believe is the first acute Phase I study in very advanced PD patients to determine if a D1 agonist can be studied safely in this population. That study (Huang et al., 2020) shows that even these very advanced patients (Hoehn & Yahr >4) can be studied effectively, and that at least one D_1_ agonist was well-tolerated. In the absence of a cure on the horizon, and with development of D_1_ agonists finally accelerating(Mailman et al., 2001), the possibility that we can bring new hope to PD patients at the bleakest part of their journey cannot be disregarded, especially because D_1_ receptors play important roles in processes like cognition (Arnsten et al., 2017). As pharmacologists, we should also lobby our neurologist colleagues to eschew the use of the general term “dopamine agonists” which currently define drugs that all prefer D_2_-like receptors (Millan et al., 2002). It is long overdue to retire this term, and replace it with a name of greater pharmacological precision as is done elsewhere in medicine.

## 5. Funding sources and competing interests

This work was supported, in part, by Public Health Service research grants MH040537 (RBM), NS105471 (RBM), and NS105471 (XH). The authors thank the Axion Research Foundation for the technical support critical for these experiments.

Dr. Mailman is an inventor of D_1_ agonist technology, the conflicts-of-interest of which are managed by the Pennsylvania State University College of Medicine. He is the past recipient of research funds and consulting compensation from Pfizer, Inc. Dr. Huang is an inventor of D_1_ agonist-related technology whose interests were assigned to the University of North Carolina. Dr. Huang has also received nominal transportation and per diem expenses from Acadia, Medtronics, and Cerevel Therapeutics, and research support from Pfizer, Biogen, and Biohaven. Drs. Mailman and Huang have no other conflicting interests. Dr. Yang has no conflicting interests. The opinions in this review are those of the authors alone and do not reflect those of the university or any other party.

## 6. Author contributions

The study was conceptualized by Dr. Mailman, who also provided the dihydrexidine and remoxipride. The experiments were conducted by the staff of the Axion Research Foundation. The data were interpreted, and the manuscript prepared and revised, by Drs. Mailman, Huang, and Yang. All authors have approved the submitted copy.

## 7. Ethics statement animal experimentation

Care and treatment of these monkeys were in compliance with the US Public Health Service Guide for the Care and Use of Animals (1985). The protocols were approved by the Axion/St. Kitts Biomedical Research Foundation animal care and use committee.

## 9. Acknowledgements

The authors offer special thanks to Professor Jane R. Taylor for extremely valuable and insightful comments about this study.

## Notes

### Competing Interest Statement

Dr. Mailman is an inventor of D1 agonist technology, the conflicts-of-interest of which are managed by the Pennsylvania State University College of Medicine. He is the past recipient of research funds and consulting compensation from Pfizer, Inc. Dr. Huang is an inventor of D1 agonist-related technology whose interests were assigned to the University of North Carolina. Dr. Huang has also received nominal transportation and per diem expenses from Acadia, Medtronics, and Cerevel Therapeutics, and research support from Pfizer, Biogen, and Biohaven. Drs. Mailman and Huang have no other conflicting interests. Dr. Yang has no conflicting interests. The opinions in this review are those of the authors alone and do not reflect those of the university or any other party.

